# Factors Influencing Emerald Ash Borer Ecological Interactions

**DOI:** 10.1101/2025.04.16.649187

**Authors:** Nicole D. Borsato, S. Eryn McFarlane, Nina Garrett, Alejandro José Biganzoli-Rangel, Daniel Marquina, Dirk Steinke, Robin Floyd, Elizabeth Clare

## Abstract

Emerald ash borer beetles (*Agrilus planipennis*) in North America are a destructive invasive species that increase tree mortality continent-wide, resulting in major ecological and economic impacts. Trees that are infested experience mortality rates which can exceed 99%, disrupting ecological communities and threatening the $218 billion forestry industry in North America. Given the ecological and economic impact of these pests, we seek to identify biological interactions and gain a better understanding of what ecological factors might influence these relationships. We use DNA metabarcoding from multiple markers to analyze the fungal, parasitic, plant, and microbial interactions of these beetles, and assess the relative importance of life stage (e.g., larvae, pupae, and adults), collection location, habitat, and date on the detection of ecological interactions. We detected 30 different taxonomic orders including 29 order-level interactions in larva1-stage individuals (3 animal, 17 bacteria, and 9 fungi), 64 in larva2-stage individuals (8 animal, 24 bacteria, and 32 fungi), 10 in larva3-stage individuals (2 animal, 3 bacteria, and 5 fungi), 74 in the pupae (5 animal, 31 bacteria, and 38 fungi), and 82 in the adult beetles (4 animal, 48 bacteria, 29 fungi, and 1 parasitic alveolate). These detections include several likely agents of biocontrol including the known commercially available *Beauveria* fungus, and several potential parasites including *Wolbachia* and ichneumonid wasps. A random forest model suggests the detection of interactions is best predicted by collection date and life stage, with interactions more likely to be detected in pupal samples which may be the ideal target for future analysis, where cost and time constraints prevent the more thorough analysis of all life stages.

## Introduction

### Emerald Ash Borers in North America

The invasive emerald ash borer (*Agrilus planipennis* Fairmaire, 1888) is a wood-boring beetle native to northeastern Asia and was likely introduced to North America from the Tianjin City/Hebei region of China (Bray et al., 2011). It was first detected in North America near Detroit, Michigan in summer 2002, although it is believed that the first individuals were introduced to the area in the 1990s and remained undetected (Cappaert et al., 2005; Siegert et al., 2014). Since its detection in 2002, the beetle has spread to five provinces in southeastern Canada (Ontario, Quebec, Manitoba, Nova Scotia, and New Brunswick) and 36 states in the United States of America, the majority of which are in northeastern regions adjacent to Canada (Natural Resources Canada, 2013; United States Department of Agriculture, 2023).

These forest pests primarily target stressed ash (*Fraxinus* spp.) trees (e.g., trees that are girdled, wounded, or growing in unfavourable conditions), although healthy trees are susceptible to beetle colonization as well (Cappaert et al., 2005; McCullough, Poland, Anulewicz, et al., 2009; McCullough, Poland, & Cappaert, 2009; X.Y. Wang et al., 2010; Wei et al., 2004). The larvae consume the phloem and cambium of the ash trees they inhabit, leaving behind characteristic S-shaped galleries in their wake (Figure 1) that disrupt the transportation of nutrients throughout the tree. Many of these galleries also score the outer sapwood (xylem), further disrupting the flow of water and nutrients in affected trees (Cappaert et al., 2005; X.Y. Wang et al., 2010). Emerald ash borer infestations are associated with severe ecological and economic impacts. Infestations are associated with extensive ash tree canopy loss, which results in tree mortality rates (including those of mature, healthy trees) often exceeding 99% within the first decade of emerald ash borers arriving in an area (Klooster et al., 2014; Knight et al., 2013; Natural Resources Canada, 2013) and, as a result, six North American ash species are now included in the International Union for Conservation of Nature’s (IUCN) Red List of Threatened Species (International Union for Conservation of Nature, 2022).

**Figure 1.**
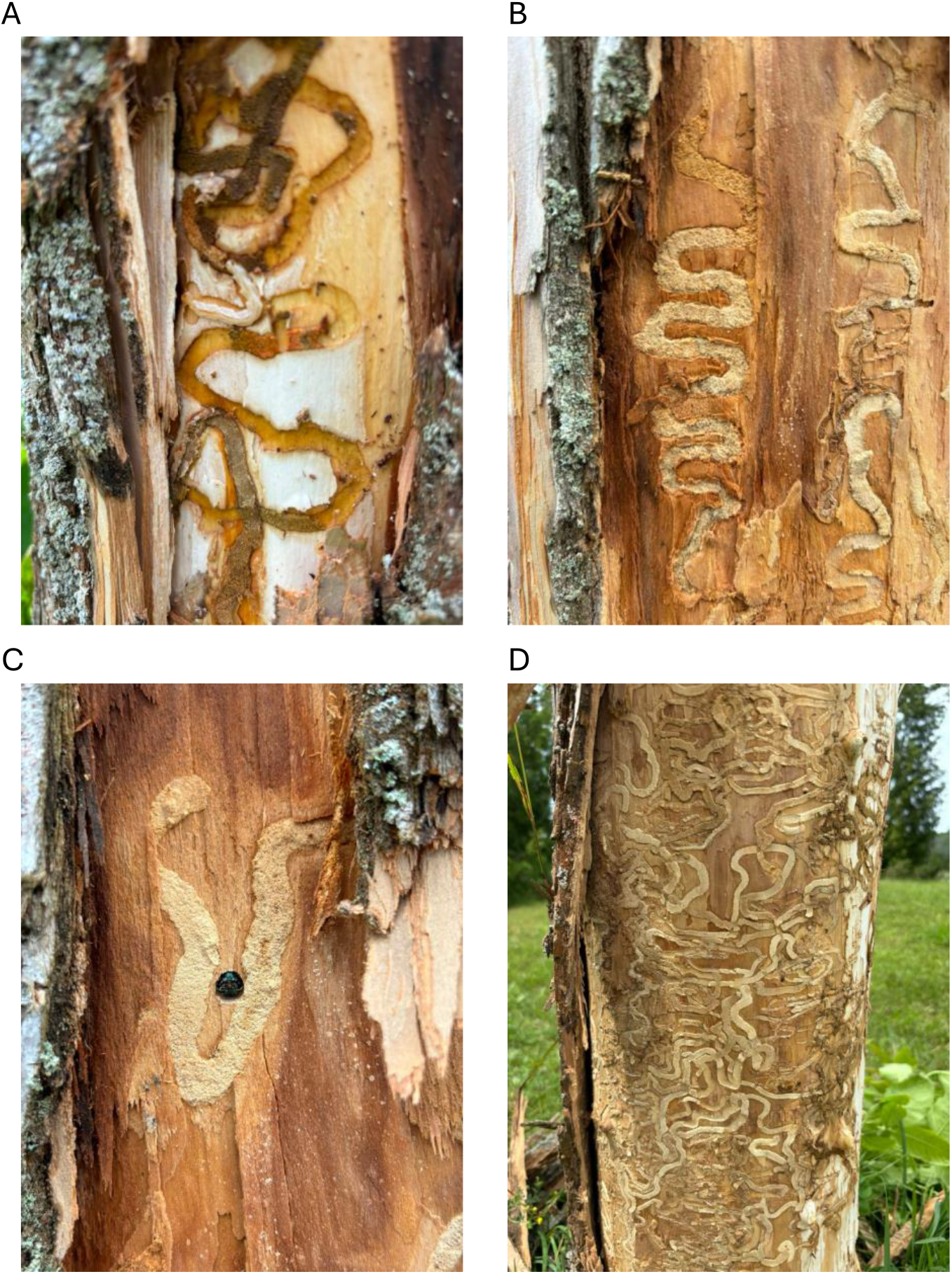
Extensive damage caused by emerald ash borers. A) a larva *in situ* cutting a trail just under the bark, B) characteristic “S” shape paths cutting off fluid and nutrient transport, C) an adult in a recessed tunnel extending through the bark and D) extensive damage just under otherwise normal looking bark of an ash tree which had lost >90% of its canopy and was scheduled for removal. All images taken on the Bruce Peninsula, Ontario Canada July 20 2024.

Increased ash tree mortality reduces the availability of food and shelter for organisms that rely on those trees (e.g., eastern ash bark beetle (*Hylesinus aculeatus*), black-headed ash sawfly (*Tethida barda*), etc.), alters the structure of ecological communities (e.g., canopy dieback creates gaps that allow light to reach subcanopy levels, encouraging understory species, such as *Acer* spp., over infested ash trees, altering the distribution and abundance of taxa; Canham, 1988; Flower et al., 2013; Gandhi & Herms, 2010), and threatens the $218 billion gross domestic product forestry industry in Canada and the United States of America (Aukema et al., 2011; Natural Resources Canada, 2022; United States Department of Agriculture, 2021). Since their introduction in the 1990s, emerald ash borers have killed tens of millions of trees (Cappaert et al., 2005) and are estimated to be the costliest forest pest in North America, responsible for approximately $850 million annually in local government expenditures (i.e., tree removal, replacement, and treatment) and approximately $380 million annually in lost residential property values in the USA alone (Aukema et al., 2011).

### Management of Emerald Ash Borer Populations

There have been multiple unsuccessful attempts to eradicate emerald ash borer populations in North America since their detection in 2002. This lack of success is partly attributed to infestations being difficult to detect until the population density increases and the host begins exhibiting external symptoms of an infestation (e.g., crown dieback, bark splits, etc.; Cappaert et al., 2005). Early suppression efforts included quarantine areas (where populations were established; discontinued by United States Department of Agriculture – Animal and Plant Health Inspection Service (USDA-APHIS) in 2021; Hope et al., 2021; Removal of Emerald Ash Borer Domestic Quarantine Regulations, 2020) and the regulation of the transportation of wood products from infested quarantine areas (both in North America and beyond; Government of Canada, 2013a; Hope et al., 2021; Removal of Emerald Ash Borer Domestic Quarantine Regulations, 2020).

Other suppression efforts make use of a combination of physical, chemical, and biological controls. However, these methods are not very effective, or come with negative consequences. Physical controls include creating ash-free firebreak zones and the removal of infested trees. However, adult beetles can fly far enough to cross firebreak zones or can be established outside the zone prior to creation without being noticed (Herms & McCullough, 2014; Taylor et al., 2010) which makes firebreak zones ineffective. Additionally, the removal of infested trees is dependent on the infestations having visible external effects on the host tree. Chemical controls include the application of insecticides (e.g., imidacloprid, dinotefuran, azadirachtin, and emamectin benzoate; Herms & McCullough, 2014; McCullough, 2020) that may have unintended consequences in the environment, which can limit their usefulness and areas they can be applied to. Biological controls include the release of natural predators and parasitoids of emerald ash borers (i.e., the parasitoid wasps *Tetrastichus planipennisi, Spathius agrili, Spathius galinae*, and *Oobius agrili*) that were imported from China and Russia (Duan et al., 2018; Government of Canada, 2013b). Species native to North America, such as woodpeckers (Picidae), parasitoid wasps (e.g., *Atanycolus cappaerti*), and the fungus *Beauveria bassiana* (now commercially available as FraxiProtec), have also been linked to decreased infestation levels (Cappaert et al., 2005; Cappaert & McCullough, 2009; Lindell et al., 2008; Srei et al., 2020) but the effectiveness of these biological controls varies, and it is unclear whether potential agents are functional in the wild. Woodpecker predation, for instance, is dependent on pest population density and the condition of the ash trees (Jennings et al., 2013; Lindell et al., 2008). Similarly, the effectiveness and horizontal transmission of *Beauveria* may depend on the density of infested trees and pest populations (Srei et al., 2020). Additionally, one of the introduced parasitoids (*S. agrili*) was found to be unable to establish self-sustaining populations in regions north of 40° N latitude and has yet to establish well in any release sites south of that latitude. However, the other three (*T. planipennisi, S. galinae*, and *O. agrili*) were successful in doing so beyond their initial release sites (Duan et al., 2023; Government of Canada, 2013b) and tracking their natural dispersal is key to understanding their effectiveness.

### Ecological Interactions in Biological Controls

Trophic interactions play an important role in regulating emerald ash borer populations (Cappaert et al., 2005; Cappaert & McCullough, 2009; Lindell et al., 2008; Srei et al., 2020). Borsato et al. (in press) compared DNA metabarcoding techniques to generate data on potential interactions from molecular traces across a variety of insect taxa. In this approach, a “host” organism is subjected to DNA extraction and PCR of target molecular markers to amplify co-occurring DNA signatures (i.e., bacteria, fungi, parasites, etc.) that interact with an individual of interest (e.g., by residing in or on a host organism). Interactions may leave behind a genetic trace signature that can be detected forensically, allowing a metabarcoding approach to be used.

Borsato et al. 2025 (in press) tested multiple target gene regions for the identification of these interactions and generated a dataset as part of methods development. The objective of this study is to use these data generated for emerald ash borers to classify the ecological factors that influence potential ecological interactions. In particular, we assess the impact of 1) life stages (e.g., larvae, pupae, and adults), and 2) ecological correlates (e.g., collection date, location, and habitat) on the detection of potential ecological interactions of emerald ash borer hosts. We also look specifically for potential agents of biological control (i.e., fungi, parasites, etc.) or particularly vulnerable pest life stages to highlight any approaches to suppress or better target this invasive species with minimum impact on ecosystem.

## Materials & Methods

The data used in this study are described in Borsato et al. (in press). In brief, emerald ash borer larvae (n= 54 size 1, n=68 size 2, n=41 size 3), pupae (n=53), and adults (imagoes; n=60) were hand collected from ash trees found in Brockville (n = 185), Ennotville (n = 41), Guelph (n = 41), and Puslinch (n = 10) in Ontario, Canada by the collection unit of the Canadian Center for Biodiversity Genomics (see Appendix B, Tables S1 and S2). The specimens were crushed inside a tube using a pestle and DNA was extracted using the Qiagen blood and tissue kit following the manufacturer’s guidelines with a decreased final DNA elution volume of 150 µl of Buffer AE to increase DNA yields. To account for differences in biomass, the DNA was extracted from the posterior half of the abdomen for imago specimens, the posterior third for pupa specimens, a 6 mm fragment of the posterior end of size 2 (larva2) and 3 larva (larva3) specimens, and the whole size 1 larva (larva1).

To confirm larva ID and identify potential insect parasites we amplified a partial COI barcode using the ZBJ-ArtF1c/ZBJ-ArtR2c primers (Zeale et al., 2011). The interal transcribed spacer (ITS) can be used to amplify both fungi and plants but different primer sets show differential amplification. Because host plant was known (ash trees) we used ITS3/ITS4 primers (White et al., 1990), which have a better recovery for fungi to amplify potential symbionts. We used the 799F-mod3/1115R primers targeting 16s (Hanshew et al., 2013; Reysenbach & Pace, 1995) to amplify microbial taxa. All PCR products were sequenced on the Illumina MiSeq v3 using a 600-cycle run at the Barts and the London Genome Center at Queen Mary University of London, Blizzard Institute (UK). To target parasites, DNA extracts were sent to the University of Guelph where a long-read 18S region was amplified using SSU_F_07/SSU_R_26 primers (Carta & Li, 2018; Floyd et al., 2002) and sequenced using the Oxford Nanopore MinION platform on a Flongle flowcell. All PCR primers, reaction mix, thermocycling conditions, and positive and negative controls are detailed in Borsato et al. 2025 (in press). Bioinformatics processing was performed using DADA 2 (ITS), mBRAVE (COI), QIIME2 (16S), and a purpose built platform for MinION Data (18S; Hebert et al., 2023). Full analysis details are given in Borsato et al. 2025 (in press).

To assess the relative influence of age, location (i.e., region and sector), habitat, sub-habitat, and collection time (day of the year) we constructed a random forest classification model (Breiman, 2001) where the presence or absence of order-level interactions was classed as a response variable, and specimen age, collection region, collection sector, collection habitat, collection sub-habitat, and collection date were treated as predictors (Appendix B, Tables S1 to S3, and Appendix C). We then considered fungi, microbes, and parasites as separate taxonomic groups and repeated the analysis using these taxa as separate response variables. The random forest models were generated using the *caret* package (Kuhn, 2008) in R Studio (R Core Team, 2023). A random forest model was selected over other models due to its robust nature (e.g., ability to handle both continuous and categorical variables and outliers), high accuracy, and ability to evaluate variable importance (i.e., which variables most influence the accuracy of the random forest model predictions; Breiman, 2001).

To ensure reproducibility, the seed was initialized to 42 using the set.seed function prior to generating the models. Data was then randomly split into “training” (70%) and “testing” (30%) sets, with the former used to train the random forest model. Repeated k-fold cross validation (k = 10, repeats = 5) was included in each model to estimate the performance (accuracy) of the models, and to improve model accuracy by allowing more data to be used when training the model. The number of variables tried in each split (mtry) was determined using the tuneRF function in the *randomForest* package (Liaw & Wiener, 2002), where mtry = 2 was found to be optimal for the animal and overall models, while mtry = 1 and mtry = 4 were found to be optimal for the bacteria and fungi models, respectively. The testing set was used to evaluate model accuracy by generating a confusion matrix using the *caret* package (Kuhn, 2008). Variable importance (i.e., the mean decrease in the Gini coefficient) was also determined using the *caret* package (Kuhn, 2008). Accumulated local effects (ALE) plots were then generated from the training data using the *ilm* package (Molnar et al., 2018) to determine the impact each variable has on the presence or absence of interactions. A correlation matrix was also generated using the *stats* (R Core Team, 2023) and *ggcorrplot* packages (Kassambara, 2023) to determine the correlation between predictor variables.

## Results

The species interaction data used here are fully described by Borsato et al. 2025 (in press). In summary, after host data was excluded, we retained 65 identifications (61 unique taxa) from 30 taxonomic orders across the four gene regions analyzed (ITS = 23; 16S = 21; COI = 4; 18S = 17). Using the COI target region four arthropod orders were detected, including those of potential parasites. Due to strict quality filtering protocols, no ASVs were retained from the larva1 samples. However, two orders were identified in the larva2 samples, one in the larva3 and pupae samples, and two in the imago samples. These identifications included a family of parasitoid wasps (Ichneumonidae, order Hymenoptera). We detected 13 orders of bacteria using the 16S marker, with larva1 specimens hosting a microbial richness of five orders, larva2 having three orders, larva3 having two orders, pupae having eight orders, and imagoes having nine orders. Notably, the insect pathogen *Wolbachia* sp. (order Rickettsiales) was detected in only one sample. The ITS region allowed the detection of 11 fungal orders, with five orders detected in larva1, seven orders in larva2, three in larva3, six in pupae, and eight in imagoes. Notably, *Beauveria* sp. (order Hypocreales), a fungal biocontrol agent, was detected in two samples. Although *Beauveria sp*. was also detected in one of the negative controls, the identification was ultimately kept due to the very low read abundance in this control, compared to the read abundance in the emerald ash borer samples, and there being no lab-based source of contamination. The detection of *Beauveria* sp. is also supported by the 18S data, where it was also detected at order-level in the same samples. We detected nine orders using the 18S region, with six of the nine representing fungi (overlapping with those identified using ITS) and the remaining three representing nematodes, insects, and a parasitic alveolate protist (*Cryptosporidium* sp., order Cryptosporida). We identified three orders in the larva1 samples, five in larvae2, one in larvae3, four in pupae, and two in imagoes.

These identifications account for 259 potential order-level interactions across emerald ash borer life stages (Figure 2). In total, we retained 29 order level interactions in the larva1 samples (three animal, 17 bacteria, and nine fungi), 64 in the larva2 samples (eight animal, 24 bacteria, and 32 fungi), ten in the larva3 samples (two animal, three bacteria, and five fungi), 74 in the pupal life stage (five animal, 31 bacteria, and 38 fungi), and 82 in the imago life stage (four animal, 48 bacteria, 29 fungi, and one parasitic alveolate). Overall, the adult life stage had the most potential interactions detected, and the majority of interactions detected occurred between emerald ash borers and fungi.

**Figure 2.**
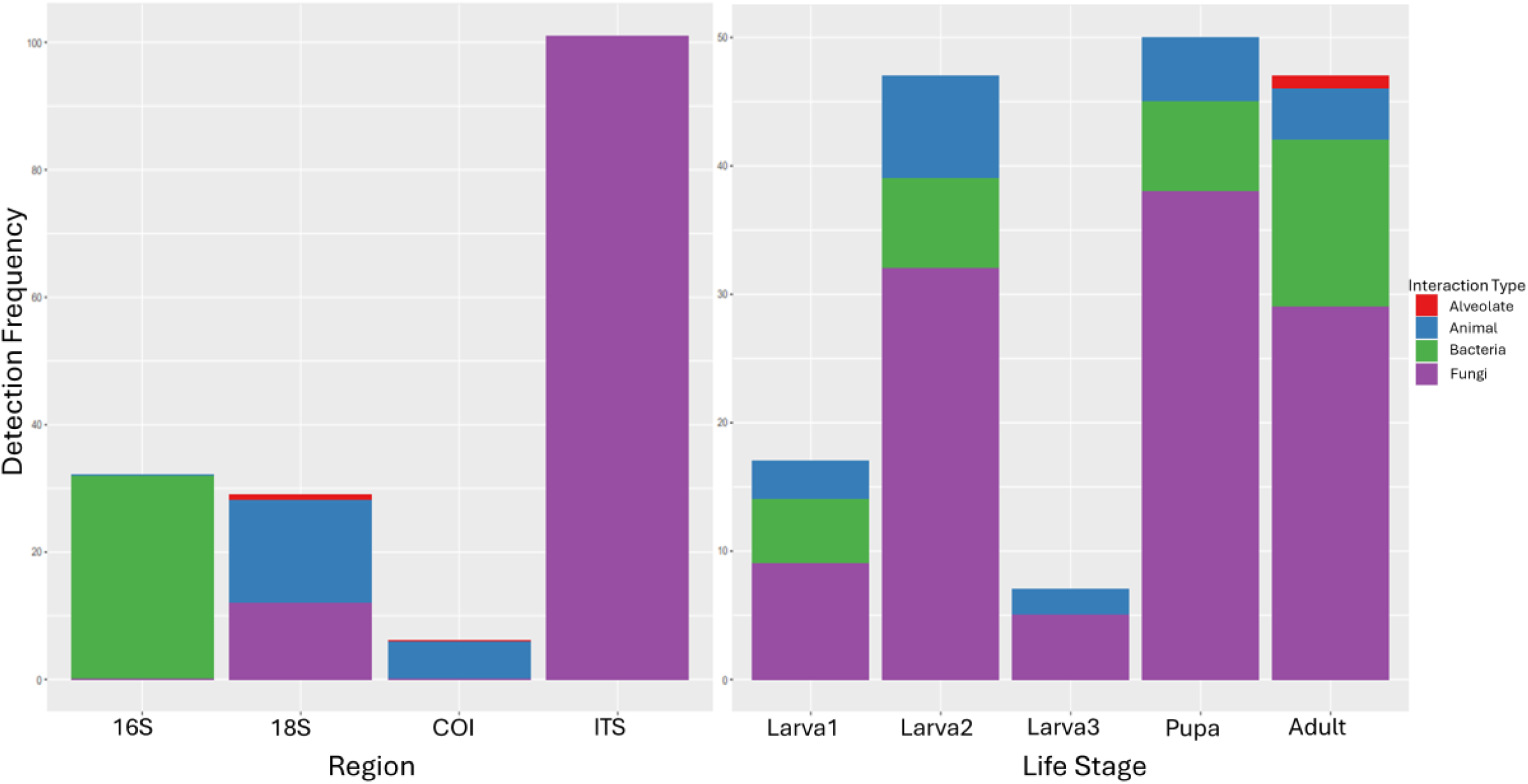
Interactions by target region and in life stages of emerald ash borers. The recovery of various targets by target amplified region (left) shows high variability in taxonomic richness. The profile of taxonomic recovery of co-amplified taxa is highly variable between life stages (right).

The classification accuracy of the total interaction random forest model was 68.92%, with an out-of-bag (OOB) error estimate of 31.18%. The model was found to correctly classify the testing data 69.23% of the time. The three most important features (Figure 3), determined by the mean decrease in the Gini coefficient (MDG), were larva3 life stage (MDG = 4.94), collection date (represented as day of the year; MDG = 3.72), and pupa life stage (MDG = 1.76). The ALE plots indicated that interactions were more likely to be present in the pupal life stage (Figure 4), in samples collected prior to July 14^th^ (195^th^ day of the year), and in samples collected from the Ennotville region, Rosedale sector, woodland habitat, and forest sub-habitat (Appendix A, Figures S1 to S5).

**Figure 3.**
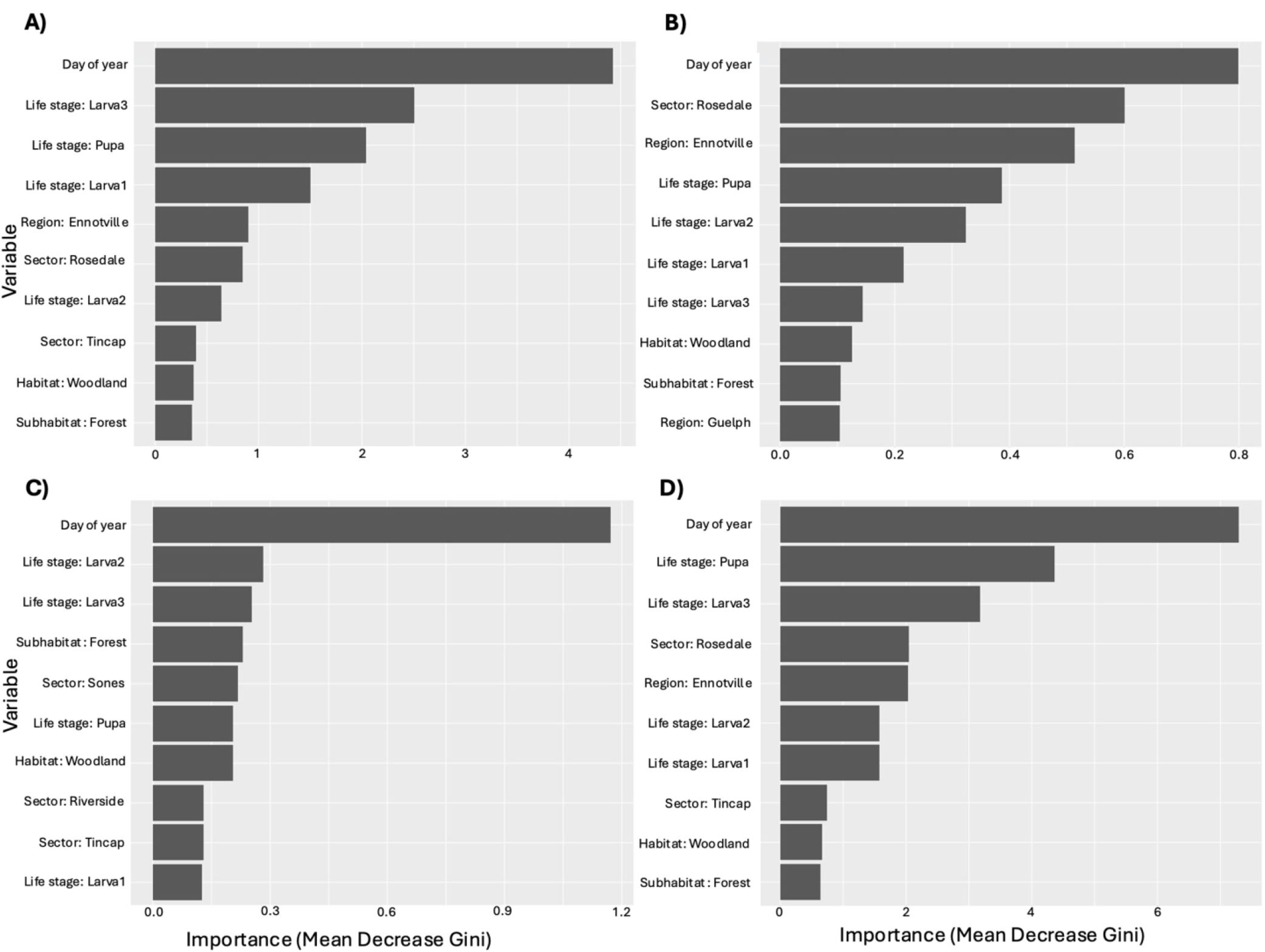
Top ten most important variables in random forest models. The ten variables with the highest importance (mean decrease Gini) are ordered top to bottom from most important to least important in classifying the presence or absence of interactions in A) the overall interaction model, B) the animal interaction model, C) the bacteria interaction model, and D) the fungi interaction model.

**Figure 4.**
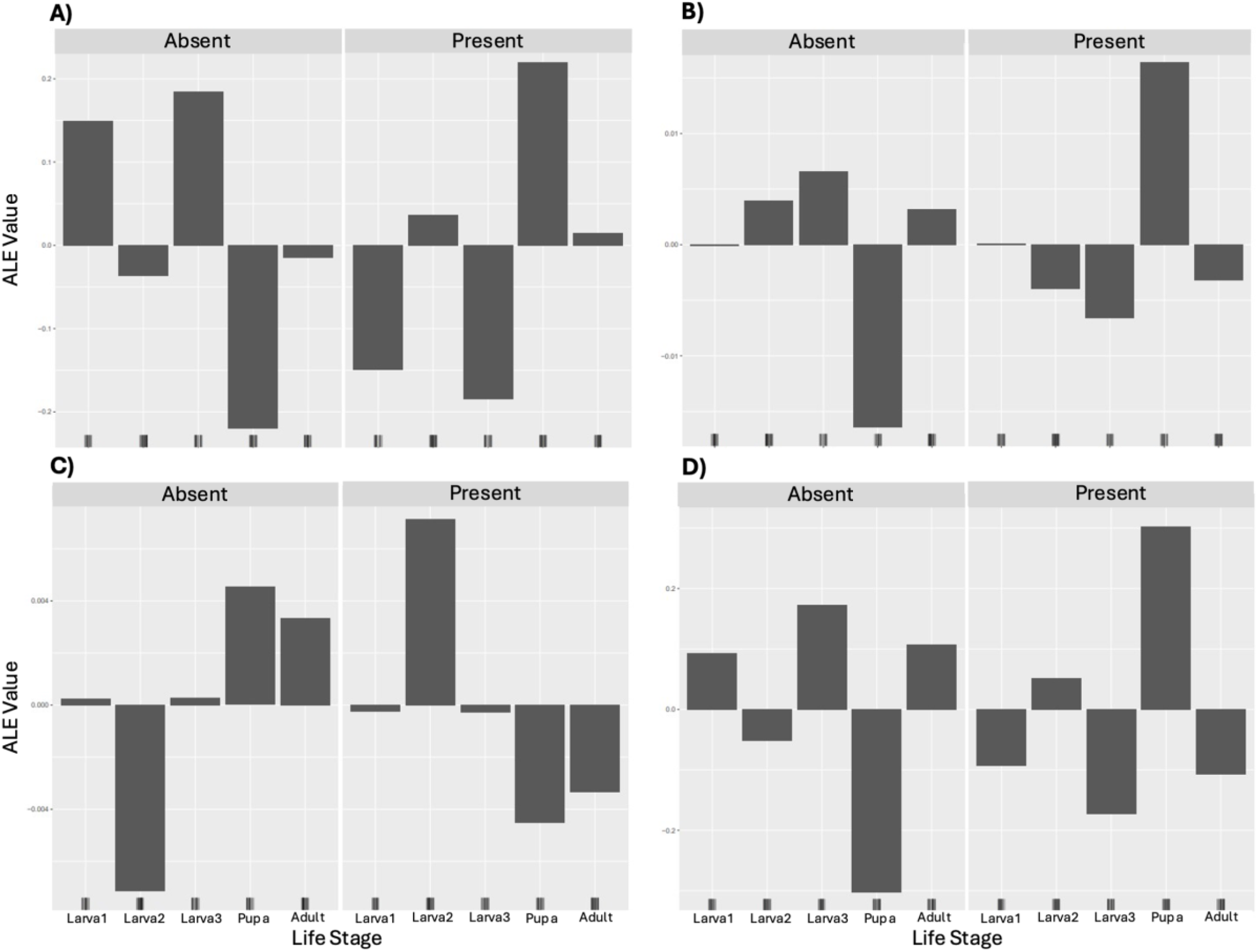
Effect of emerald ash borer life stage on the detection of interactions. Accumulated Local Effects (ALE) plots portray the effect life stage had on the presence or absence of A) overall interactions, B) animal interactions, C) bacteria interactions, and D) fungi interactions

For bacterial interactions, the estimated classification accuracy of the random forest model was 68.28%. The model had an OOB error estimate of 31.72% and it was found to correctly classify the testing data 69.23% of the time. The three most important features (Figure 3) were collection date (MDG = 1.64), larva3 life stage (MDG = 0.17), and Tincap sector (MDG = 0.54). The ALE plots indicated that interactions with bacteria were more likely to be present in the pupal life stage (Figure 4), in samples collected prior to June 22^nd^ (173^rd^ day of the year), and in samples collected from the Guelph region, Sones sector, woodland habitat, and forest sub-habitat (Appendix A, Figures S1 to S5).

The fungal interactions random forest model had an estimated classification accuracy of 68.81% and an OOB error estimate of 31.18%. The fungi random forest correctly classified the testing data 76.92% of the time, and the three most important variables (Figure 3) were collection date (MDG = 7.28), pupa life stage (MDG = 4.36), and larva3 life stage (MDG = 3.17). The ALE plots indicated that interactions with fungi were more likely to be present in the pupal life stage (Figure 4), in samples collected prior to July 6^th^ (187^th^ day of the year), and in samples collected from the Ennotville region, Rosedale sector, artificial habitat, and urban sub-habitat (Appendix A, Figures S1 to S5).

The estimated classification accuracy of the animal interactions random forest model was 92.49%. The OOB error estimate for the model was 7.57%. It was found to correctly classify the testing data 92.41% of the time. The three most important variables (Figure 3) were collection date (MDG = 0.80), Rosedale sector (MDG = 0.60), and Ennotville region (MDG = 0.51). The ALE plots indicated that interactions with animals (i.e., insects, nematodes, etc.) were more likely to be present in the pupa life stage (Figure 4), in samples collected after July 21^st^ (202^nd^ day of the year), and in samples collected from the Ennotville region, Rosedale sector, artificial habitat, and arable sub-habitat (Appendix A, Figures S1 to S5). The correlation between most of the predictor variables was low, although there was high correlation detected between some variables (e.g., the correlation between the Rosedale collection sector and Ennotville collection region was high, which was unsurprising given Rosedale is a sublocation of Ennotville; Appendix A, Figure S6).

## Discussion

Ecological interactions play an important role in the evolution and survival of a species (Barraclough, 2015; Thompson, 1999). In the case of emerald ash borers, interactions with biological controls (e.g., woodpeckers, *Beauveria*, etc.) may lead to population control of the pest. We applied data generated by metabarcoding of four genetic markers to assess the factors which influence the potential ecological interactions across the tree of life of the destructive (in North America) emerald ash borers. Our objectives were to determine whether different life stages and ecological correlates can predict the detection of non-target taxa within a sample. Our model suggests that life stage targeted, collection date, and location (region and sector) can predict interactions, while habitat and sub-habitat play minor roles (Figure 3). We identified interactions with a variety of taxa (e.g., bacteria, mites, fungi, nematodes, etc.), including interesting potential agents of biocontrol (e.g., *Beauveria* sp., parasitoid wasps) in several samples of emerald ash borer.

### Taxonomic Identifications

A total of 259 potential order-level interactions were detected across emerald ash borer life stages. These interactions represent 61 unique taxa from 30 taxonomic orders across the four markers used in this analysis (COI, ITS, 16S, and 18S). Excluding host data from the COI data, we identified four orders of arthropods, which included one family of gall midge (Cecidomyiidae, order Diptera), two families of mites (Oppiidae, order Sarcoptiformes, and Eupodidae, order Trombidiformes), and notably, ichneumonid parasitoid wasps (order Hymenoptera, see biological control discussion below). The 16S marker was used to detect bacterial interactions, although the majority of the bacteria associated with our samples are ubiquitous throughout the environment and can be associated with soil (e.g., *Terriglobus* sp.) or the rhizosphere (e.g., *Neorhizobium* sp.). Many of these detections are thus likely co-occurring interactions rather than direct interactions (co-occurring and co-evolving), characteristic of the symbiome. Notably, we did detect the intracellular parasite *Wolbachia sp*. (order Rickettsiales) in one sample (see biological control discussion below). Similar to the bacteria detected by 16S, the majority of the fungi sequenced using the ITS marker are associated with soil (e.g., *Fusarium* sp.), decaying plant matter (e.g., *Yamadazyma* sp.), lichens (e.g., *Candelaria* sp.), or plant pathogens (e.g., *Diaporthe* sp.), and not the emerald ash borers themselves. It is likely that these are fungi associated with the host tree or surrounding environment, and would be classed as co-occurring rather than direct interactions. However, the fungus *Beauveria* sp. (order Hypocreales) was detected, and likely represents a true ecological interaction (see biological control discussion below). The 18S sequence data contained the greatest diversity of interaction types, including animals (i.e., nematodes and arthropods), a parasitic alveolate, and fungi. The fungal orders detected using the ITS marker overlapped with those identified using the 18S marker, even though the taxonomic resolution was the same or worse (i.e., taxa identified in both regions such as *Diaporthe sp*. and *Niesslia sp*. were only resolved at a higher taxonomic level in 18S) likely due to a lack sequence variability at lower taxonomic levels in very conserved marker regions. We also detected non-host arthropods belonging to the order Hymenoptera with the 18S marker. Detected interactions with nematodes (order Rhabditida) included Neotylenchidae (whose members can be parasitic, e.g., *Deladenus proximus*; Zieman et al., 2015), *Panagrellus* sp. (free-living nematodes associated with many habitats, including insect frass; Ferris, 2009; Srinivasan et al., 2013), and *Rhabditolaimus* sp. (bacteriophagous and commensal with other beetles, e.g., *Scolytus multistriatus*; Ryss & Polyanina, 2022). The parasitic alveolate (*Cryptosporidium* sp.) that was detected has been reported to infect humans and animals (including insects, Helmy & Hafez, 2022), thus this may be a true parasitic interaction.

### Detection of Biological Controls and Parasites

Several potential agents of biocontrol were identified across all markers used. Ichneumonidae parasitoid wasps (order Hymenoptera) were detected in larva2, larva3, and pupae samples using the COI marker. Ichneumonids have previously been observed to parasitize emerald ash borers (Duan et al., 2009), so these are likely cases of true parasitism. Additionally, the entomopathogenic fungus *Beauveria* sp. (order Hypocreales), that is known to successfully infect and neutralize emerald ash borers, was identified in two samples (one size 2 larva and one adult beetle) using the ITS marker. Although naturally occurring in the environment (e.g., in soil; Rehner et al., 2011), biocontrol products containing *Beauveria* such as FraxiProtec (GDG Environment, Canada), are commercially available and are able to significantly reduce emerald ash borer populations (Srei et al., 2020).

Using the 16S region, we identified the intracellular parasite *Wolbachia* sp. (order Rickettsiales) in one pupal sample. *Wolbachia* can infect many insects, and infections can be vertically and horizontally transmitted between hosts (Werren, 1997). Infection is associated with reproductive abnormalities, such as the feminization of genetic males, parthenogenesis induction, and reproductive incompatibility (i.e., through cytoplasmic incompatibility between hosts infected by different strains or between uninfected/infected hosts; reviewed in Werren, 1997). However it may also have a protective benefit in some viral infections (Cogni et al., 2021; Pimentel et al., 2020). The detection of *Wolbachia* is notable as it may have potential biocontrol applications (Werren, 1997; Zabalou et al., 2004).

The parasitic alveolate and nematodes identified using the 18S marker may potentially act as biocontrols as well. The alveolate (*Cryptosporidium* sp.) is known to infect insects (Helmy & Hafez, 2022), and can therefore be considered a true parasitic interaction. Similarly, members of the Neotylenchidae family (order Rhabditida) of nematodes detected can be parasitic (e.g., *Deladenus proximus*; Zieman et al., 2015), and thus this may also represent a true parasitic interaction that may have potential biocontrol applications. The detection of all these interactions suggests that some natural biocontrol may be occurring in the collection location.

### Factors Influencing the Interactions Assessed Using Random Forest Models

Four random forest models were generated in total: one to predict the presence of any interaction at all, and the other three to specifically test predictions about the presence of fungi, microbial, or animal interactions. The models were successfully able to predict the number of interactions detected using the testing dataset in a range of 69 to 92% of the time. Animal interactions were most likely to be classified correctly (∼92% of the time), while bacteria, fungal, and total interactions were less likely to be classified correctly (∼69, ∼77, and ∼69% of the time, respectively). This means that these models can be used to predict which specimens are likely to have detectable interactions based on factors such as life stage, collection date, location, and habitat up to 92% of the time. They can thus be used to identify ideal target specimens when resources limit the more thorough analysis of all specimens. In all models, collection date (represented as day of year) and life stage played the most important roles in correctly classifying the presence or absence of interactions, with the exception of the animal and bacteria models, where location (sector and region) also played a major role. It should be noted that this observed variable importance may be a result of the tendency of random forest classification models to be biased towards categorical variables with more levels (i.e., life stage and sector, which each have five levels; Strobl et al., 2007). However, features were not omitted from the models based on variable importance, so this likely does not impact the accuracy of the models used.

The ALE plots generated for each interaction type were used to determine what effect, if any, each variable has on the presence or absence of interactions detected. On our models all types of interactions were more likely to be present at the pupal life stage. Collection date also seemed to play a significant role, with interactions more likely to be present in samples collected by the end of July. This differed slightly for specific interaction types (e.g., bacterial interactions were most likely to be present in samples collected by June 22^nd^, 2022), but the latest the date ranges to is July 21^st^, 2022 (for animal interactions) before the detection rate decreases. The presence of total, fungi, and animal interactions were highest in the Ennotville collection region and Rosedale collection sector, while the presence of bacterial interactions was greatest in the Guelph region and Sones sector. Similarly, fungi and animal interactions were more likely to be present in artificial habitats, while total and bacterial interactions were more likely to be present in forest and woodland habitats. Finally, we found that bacteria and total interactions were more likely to be present in forest subhabitats, fungi interactions in urban subhabitats, and animal interactions in arable subhabitats. Future studies can therefore target specific life stages, locations, habitats, and collection dates depending on the target interactions of interest (e.g., a study that wishes to focus on fungi-beetle interactions should consider prioritizing pupae samples collected by July from artificial/urban habitats located in Ennotville).

### Novel Challenges in Differentiating interactions vs. co-occurrence

One of the main challenges in determining a true interaction from co-occurrence data is the growing understanding that environmental DNA (eDNA) likely contaminates all samples collected from the wild to some degree. eDNA includes all biological material (cells, excreta, etc.) shed by organisms continually, and it accumulates in the environment and can be collected from air (Clare et al., 2021, 2022), water (Uchida et al., 2020), soil (Foucher et al., 2020; Marquina et al., 2019), and surfaces such as leaves (Macher et al., 2023; Valentin et al., 2020) and the bodies of other animals through casual contact. eDNA accumulates and settles on biotic and abiotic surfaces where it then degrades due to environmental conditions. Consequently, eDNA of nearby organisms may be found on the surface of arthropods even if there was no direct interaction between the two. For example, Huszarik et al. (2023), demonstrated that eDNA of an exotic insect could be detected on spiders who occupied the same wet pitfall trap. We must thus expect that a sample or target organisms could become coated in eDNA simply from the environment or during collection and storage (e.g., pitfall traps, ethanol, etc.; Huszarik et al., 2023; Shokralla et al., 2010) and the duration and magnitude of this effect in wild data is only starting to be realized and little quantification of the effect has been provided. Although the ubiquitous nature of eDNA allows for its use in biodiversity and community composition assessments (Clare et al., 2022; Littlefair et al., 2023; Macher et al., 2023), it can complicate metabarcoding for species interaction quantifications, making it difficult to determine true interactions from co-occurrence. While we might speculate that true interactions leave a stronger signal, this has not been tested, and variation in recovery and amplification due to lab biases (such as primer affinity) might alter these natural occurrences.

The effect is suspected here. For example, most of the fungi detected are ubiquitous throughout the environment and are often found in soil (e.g., *Fusarium* sp.) or plants (e.g., *Alternaria* sp.). The fungus *Candelaria* sp. is known to form lichen complexes (Sanders & Masumoto, 2021) but was identified in the emerald ash borer samples despite likely not interacting directly with the specimens. Similarly, beetles and a family of gall midges (Cecidomyiidae, order Diptera) detected in the COI data are unlikely to directly interact, but the DNA was co-occurring in the same samples. Thus, many identifications are thought to reflect simple co-occurrence in the local environment. As a result, we refer to all these data as “potential” interactions in this manuscript reflecting this uncertainty.

While it is extremely difficult to envision an immediate solution to this problem, there are mitigation steps and data analysis steps which might reduce the effect. First, there is some evidence that specimens can be washed (e.g., in a bleach solution) prior to DNA extraction to remove eDNA (Binetruy et al., 2019; Hausmann et al., 2021; Huszarik et al., 2023), though this would then increase the false negative rate by removing fungi and microbes associated with that surface ecology. Dissection so that only the gut undergoes DNA extraction would allow direct interactions related to feeding and gut parasitism to be determined, and potentially increase the target DNA concentration relative to host DNA, a problem when using generalist arthropod primers when the host and parasite are both arthropods, but this would limit the scope of the interactions being assessed. However, this is very time-consuming and requires destruction of the specimen, a process objected to when analysis involves mass screening of samples or museum and collection specimens, respectively. Just as important is ecological assessment of all recovered taxa to determine if they are “reasonable” given known life history. While this will limit discovery of novel interactions, it should be encouraged so that all identifications are not automatically accepted as a true interaction.

## Conclusions

In this analysis we describe taxa co-amplified in emerald ash borer samples at different life stages and evaluate whether the detection of these potential ecological interactions can be predicted by collection date, location, life stage, or habitat using random forest models. Our models suggest that life stage is an important predictor of interactions with pupal life stage showing more interactions. Future work must consider life stage when targeting interactions, particularly when cost and time constraints limit options for analysis. We also found some support for sampling earlier in the year and regional variation in data, though with smaller sample sizes this should be treated with caution. Our data also highlights the potential environmental contamination of specimens in ecological analyses, and the difficulty in distinguishing true interactions, a challenge only starting to be considered in the vast literature on the use of DNA to identify ecological interactions. The risk of generating false positive “interactions” from contact environmental DNA requires creative solutions to estimate and mitigate the challenge.

## Acknowledgements

We would like to thank Sean Prosser at the Centre for Biodiversity Genomics for bioinformatics support and the collection unit who supported this research, particularly Gergin Blagoev, Simonne Clout, Isaiah Dowling, Hillary Hale, Reid Harrop, Paul Hebert, Chris Ho, Nao Ito, Meredith Miller, and Emily Perry.

This work was supported by The Natural Sciences and Engineering Research Council of Canada through the Discovery Grants Program, The Government of Canada’s New Frontiers in Research Fund (NFRFT-2020-0073) Tracing the Patterns of Life on a Changing Planet, Genome Canada (BIOSCAN Canada) and Ontario Genomics (OGI-208).

