## Appendix for "Factors Influencing Emerald Ash Borer Ecological Interactions"

### Appendix A: Supplemental Figures

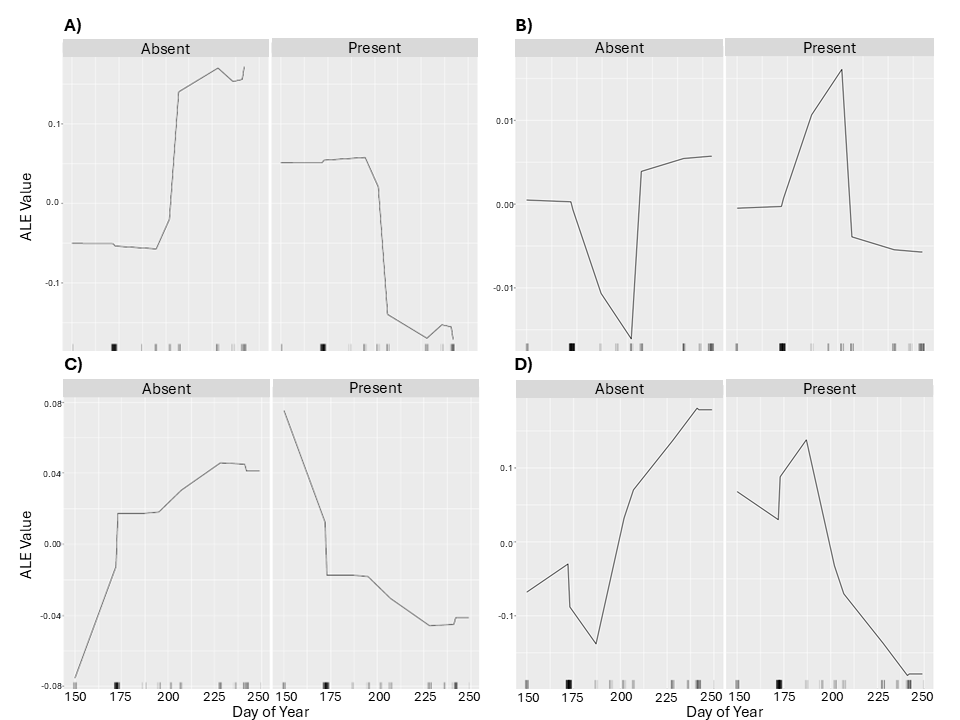

**Figure S1. Effect of emerald ash borer collection date on the detection of interactions.** Accumulated Local Effects (ALE) plots portray the effect collection date had on the presence or absence of A) overall interactions, B) animal interactions, C) bacteria interactions, and D) fungi interactions.

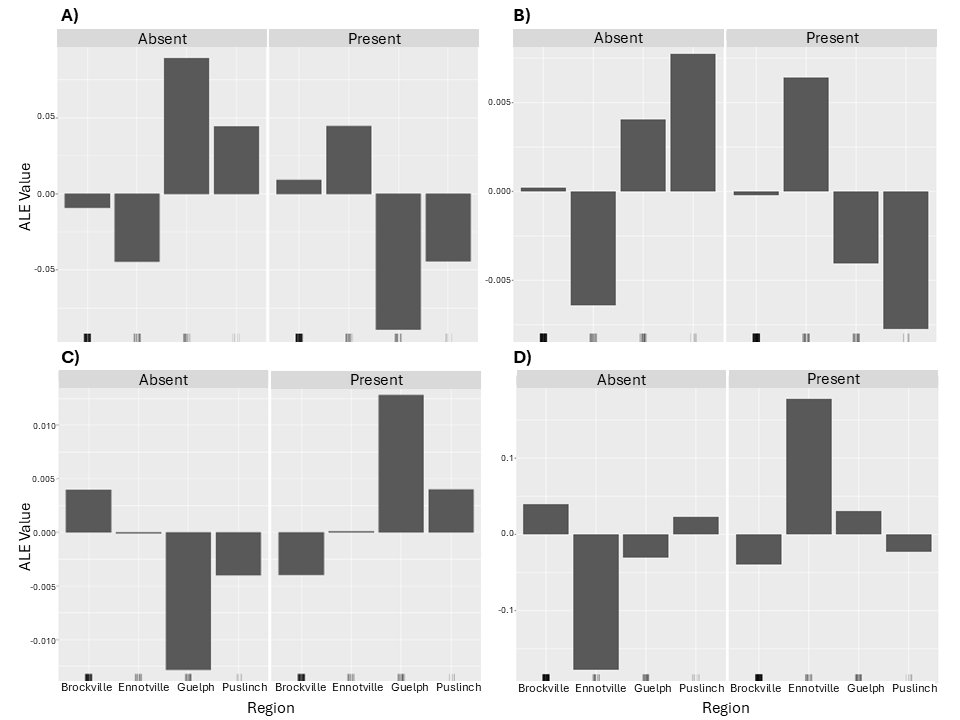

**Figure S2. Effect of emerald ash borer collection region on the detection of interactions.** Accumulated Local Effects (ALE) plots portray the effect collection region had on the presence or absence of A) overall interactions, B) animal interactions, C) bacteria interactions, and D) fungi interactions.

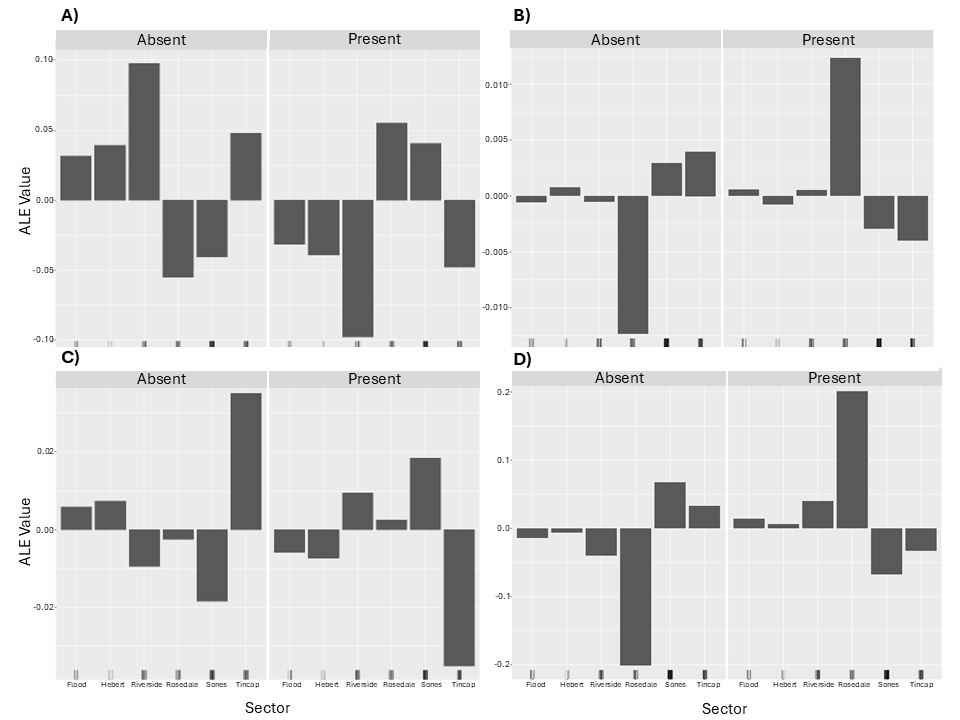
 **Figure S3. Effect of emerald ash borer collection sector on the detection of interactions.** Accumulated Local Effects (ALE) plots portray the effect collection sector had on the presence or absence of A) overall interactions, B) animal interactions, C) bacteria interactions, and D) fungi interactions.

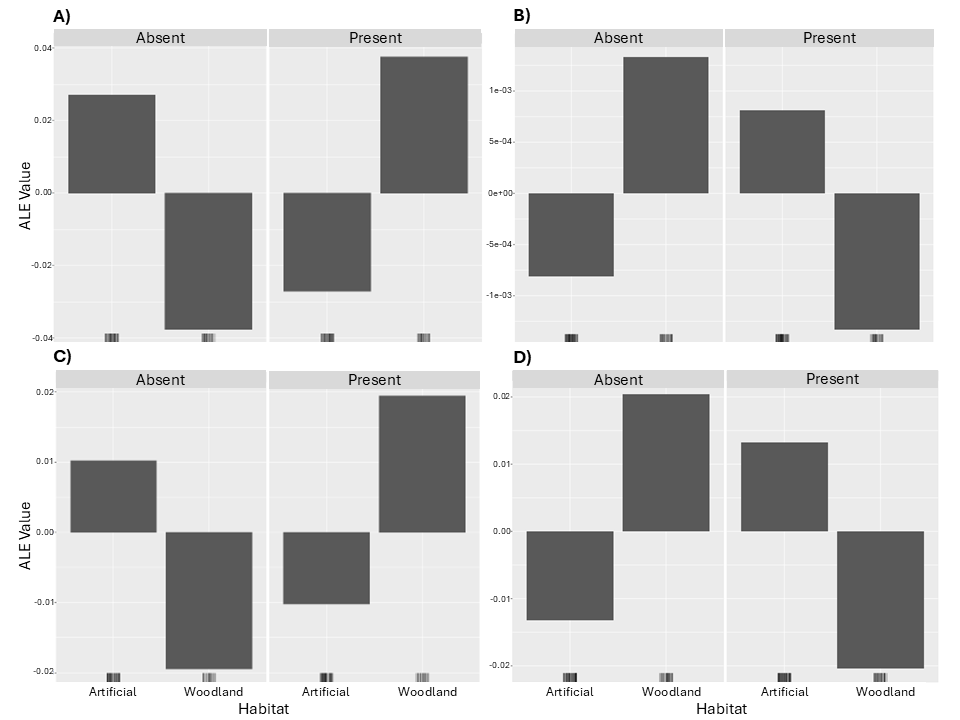

**Figure S4. Effect of emerald ash borer collection habitat on the detection of interactions.** Accumulated Local Effects (ALE) plots portray the effect collection habitat had on the presence or absence of A) overall interactions, B) animal interactions, C) bacteria interactions, and D) fungi interactions.

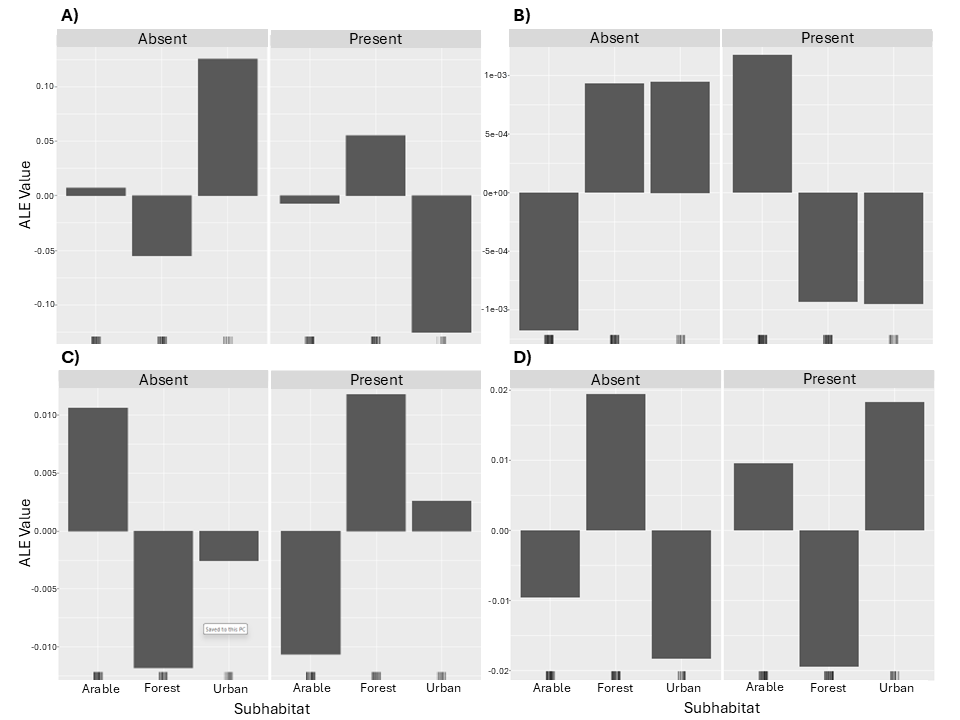

**Figure S5. Effect of emerald ash borer collection subhabitat on the detection of interactions.** Accumulated Local Effects (ALE) plots portray the effect collection subhabitat had on the presence or absence of A) overall interactions, B) animal interactions, C) bacteria interactions, and D) fungi interactions.

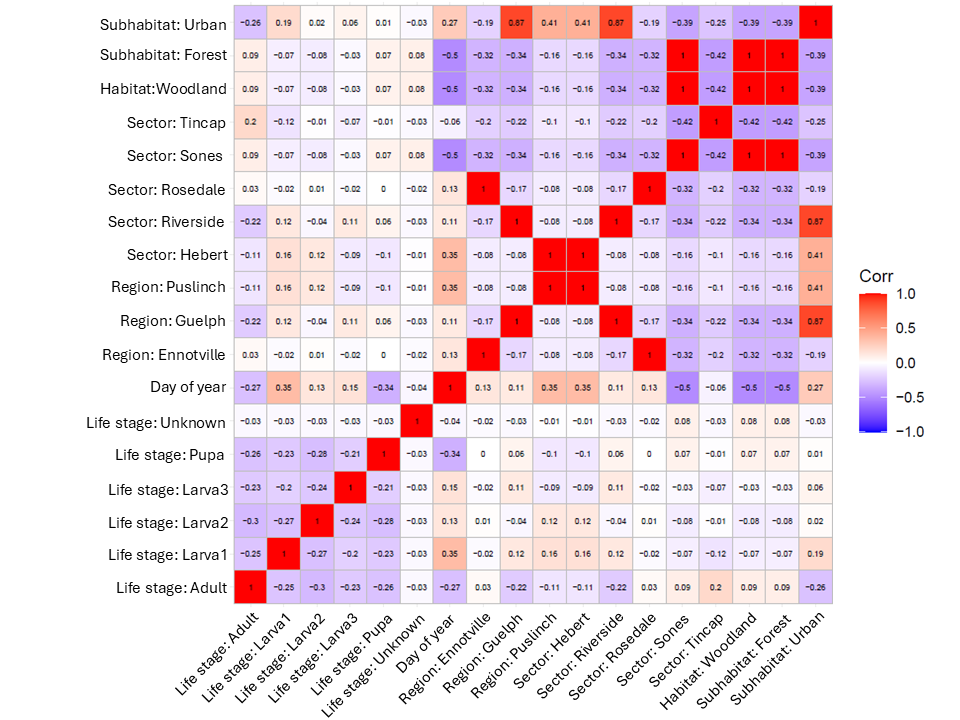

**Figure S6. Correlation matrix of the predictor variables used in the random forest models.** The correlation matrix portrays the correlation between life stage, collection date, collection location (region and sector), collection habitat, and collection subhabitat.

### Appendix B: Supplemental Tables

**Table S**1**. Emerald ash borer life stages.** A total of 277 emerald ash borer specimens were collected, including one specimen of an unknown life stage (not shown).

| Life Stage | Tissue Sampled | Specimen Count |
| --- | --- | --- |
| Larva1 (<0.5in) | Whole specimen | 54 |
| Larva2 (0.5-1in) | ~6mm fragment of posterior | 68 |
| Larva3 (>1in) | ~6mm fragment of posterior | 41 |
| Pupa | Posterior third | 53 |
| Adult | Posterior half of the abdomen | 60 |

**Table S**2**. Emerald ash borer collection locations.** A total of 277 emerald ash borer specimens were collected across southern Ontario, Canada.

| Region | Sector | Specimen Count |
| --- | --- | --- |
| Brockville | Flood | 20 |
|  | Sones | 107 |
|  | Tincap | 58 |
| Ennotville | Rosedale | 41 |
| Guelph | Riverside | 41 |
| Puslinch | Hebert | 10 |

**Table S**3**. Emerald ash borer collection habitats.** Emerald ash borer specimens were collected from a variety of habitats and subhabitats in southern Ontario, Canada.

| Habitat | Subhabitat | Specimen Count |
| --- | --- | --- |
| Artificial | Arable | 119 |
|  | Urban | 51 |
| Woodland | Forest | 107 |

### Appendix C: R Code for Random Forest Models

##Load required packages

library(randomForest)

library(caret)

library(ggplot2)

library(dplyr)

library(iml)

##Data

#load data

Fungi_Interactions <- read.csv("EAB_fungi_pa.csv")

#rename response levels

Fungi_Interactions$Fungi_Interactions[Fungi_Interactions$Fungi_Interactions==1] <- "Present"

Fungi_Interactions$Fungi_Interactions[Fungi_Interactions$Fungi_Interactions==0] <- "Absent"

#convert variable to factors

Fungi_Interactions <- within(Fungi_Interactions, {

Fungi_Interactions <- as.factor(Fungi_Interactions)

Lifestage <- as.factor(Lifestage)

Habitat <- as.factor(Habitat)

Habitat_lvl2 <- as.factor(Habitat_lvl2)

Region <- as.factor(Region)

Sector <- as.factor(Sector)

})

#view data

summary(Fungi_Interactions)

##Make the random forest model

set.seed(42) #used for reproducibility

#define repeated k fold cross validation (arbitrary values)

repeat_cv <- trainControl(method = "repeatedcv", number = 10, repeats = 5)

#create training (70%) and testing (30%) sets

index <- createDataPartition(Fungi_Interactions$Fungi_Interactions, p = 0.7, list = FALSE)

train = Fungi_Interactions[index,]

test = Fungi_Interactions[-index,]

#tune mtry value of random forest if needed, otherwise omit this line

tuning = expand.grid(mtry = 4)

rf <- train(

Fungi_Interactions~.,

data = train,

method = "rf",

trControl = repeat_cv,

tuneGrid = tuning, #remove this line if tuning is not needed

metric = "Accuracy")

print(rf)

rf$finalModel

##Tune model

tuneRF(

x = train[,1:6],

y = train[,7],

stepFactor = 0.5,

plot = TRUE,

ntreeTry = 500,

trace = TRUE,

improve = 0.05)

#optimal mtry value is 4; model was adjusted above using tuneGrid

##Evaluate accuracy of model using test data

test_prediction <- predict(rf, test[,-7])

confusionMatrix(test_prediction, test$Fungi_Interactions)

##Evaluate and plot variable importance

var_imp <- varImp(rf, scale=FALSE)$importance

var_imp <- data.frame(variables=row.names(var_imp), importance=var_imp$Overall)

var_imp%>%

arrange(importance)%>%

top_n(10)%>% #only want top 10

ggplot(aes(x=reorder(variables, importance), y=importance)) +

geom_bar(stat='identity') +

coord_flip() +

xlab('Variables') +

ylab("Importance (Mean Decrease in Gini Coefficient)")+

labs(title='Top 10 Most Important Variables')

importance(rf$finalModel) #provides importance values

##Create ALE plots

pred <- Predictor$new(rf, data = train[,-7], y = train[,7], type = "prob")

plot(FeatureEffects$new(pred, feature = "Lifestage")) +

scale_x_discrete(limits = c("Larva1", "Larva2", "Larva3", "Pupa", "Adult"))

plot(FeatureEffects$new(pred, feature = "DOY"))

plot(FeatureEffects$new(pred, feature = "Region"))+

scale_x_discrete(limits = c("Brockville", "Ennotville", "Guelph", "Puslinch"))

plot(FeatureEffects$new(pred, feature = "Sector"))+

scale_x_discrete(limits = c("Flood", "Hebert", "Riverside", "Rosedale", "Sones", "Tincap"))

plot(FeatureEffects$new(pred, feature = "Habitat"))+

scale_x_discrete(limits = c("Artificial", "Woodland"))

plot(FeatureEffects$new(pred, feature = "Habitat_lvl2"))+

scale_x_discrete(limits = c("Arable", "Forest", "Urban"))

#correlation plot

library(ggcorrplot)

model.matrix(~0+., data=Fungi_Interactions[,1:6])%>%

cor(use="pairwise.complete.obs")%>%

ggcorrplot(show.diag=TRUE, lab=TRUE, lab_size=2.5)
